# A video-based behavioural intervention associated with improved HPV knowledge and intention to vaccinate

**DOI:** 10.1101/2021.10.13.464199

**Authors:** Sarah Marshall, Anne Moore, Aoife Fleming, Laura J Sahm

## Abstract

**Aims:** The aim of this study was to design, develop and evaluate a theory and evidence-based intervention to improve human papillomavirus (HPV) and HPV vaccine knowledge, and intention to vaccinate, among parent-daughter dyads.

**Methods:** A theory and evidence-based online behavioural intervention, *“Is the HPV vaccine for me?”* was developed to improve HPV and HPV vaccine knowledge, and intention to vaccinate. The impact and feasibility of the intervention was evaluated in a prospective randomised controlled feasibility trial.

**Results:** A total of 49 parent-daughter dyads completed baseline knowledge assessment (n=24 control, n=25 intervention), and 35 dyads completed knowledge assessment at week 2 (n=17 control, n=18 intervention). The intervention was associated with a statistically significant increase in HPV, and HPV vaccine knowledge and intention to vaccinate. All intervention participants found the video interesting, while 96% found it useful.

**Conclusions:** This intervention was found to be affordable, practicable, effective (cost-effective), acceptable, safe, and equitable, in this feasibility study.

## Introduction

HPV is responsible for approximately 4.5% of global cancer disease burden, with cervical cancer the most common cancer caused by HPV infection (1). Three vaccines are licensed and marketed for use to prevent HPV infections and their sequelae: Cervarix^®^, Gardasil^®^ and Gardasil 9^®^. These are intended to be administered to adolescents before the onset of sexual activity (1). The safety of these vaccines is well established (2). However, unsubstantiated claims linking the administration of these vaccines to the development of a plethora of adverse effects (1), has led to a significant reduction in vaccine uptake worldwide (3). There is therefore a need to develop interventions to support positive vaccine decision-making (4). Several interventions have been designed to address HPV vaccine hesitancy, frequently targeting parents (5), and evaluation is often based on the impact of the intervention on parents’ intention to vaccinate, with several studies reporting a statistically significant impact on intentions (6, 7). However, it has been recommended that adolescents be included in healthcare decision-making (8). Therefore, in this study, the target population of our behavioural intervention is parent-daughter dyads.

Interventions have been shown to be more effective if they are based on principles drawn from evidence and theories of behaviour change (9). Evidence was generated in a comprehensive systematic review (10), and through a series of qualitative research studies, in both female adolescents (11) and parents. These qualitative studies were guided by behaviour change theory, including the Theoretical Domains Framework (TDF) (12), the COM-B model (13), the Behaviour Change Wheel (BCW) (13), and the Behaviour Change Technique Taxonomy version 1 (BCTTv1) (14). The TDF guided the development of focus group and interview topic guides, and was used as a coding framework in data analysis (13). Ten of the 14 TDF domains were selected as the most relevant (12, 15) (Table 1).

**Table 1.**
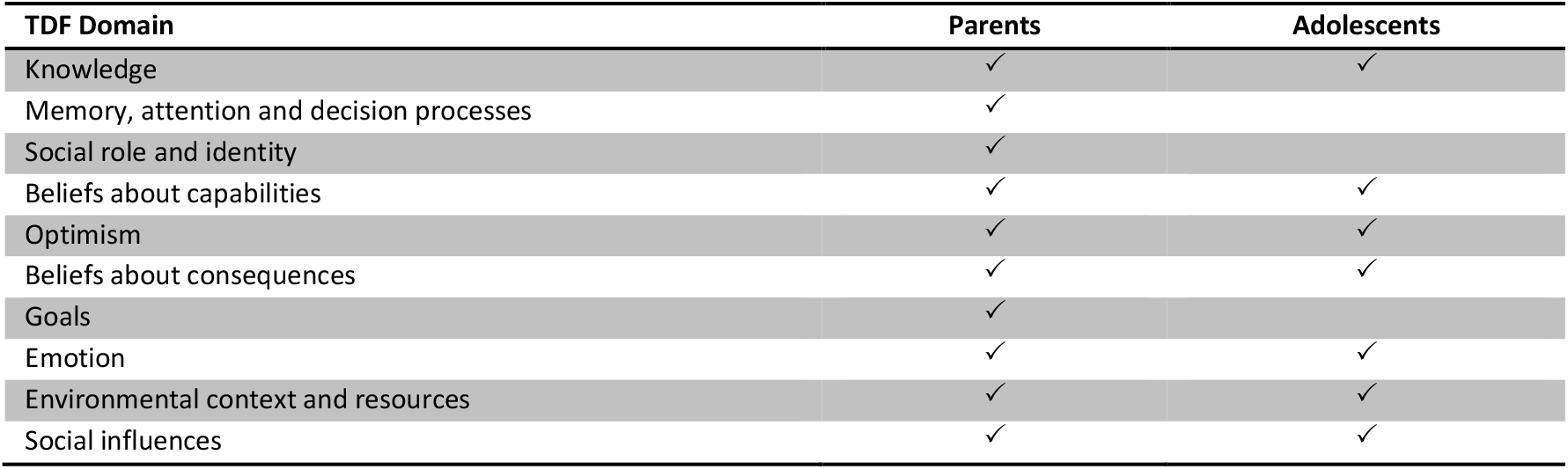
Theoretical Domains Framework (TDF) Domains identified through qualitative research (11)

The ten TDF domains were linked to several components of the COM-B model: psychological capability; physical and social opportunity; and reflective and automatic motivation. The BCW was then used to identify five relevant intervention functions: education; persuasion; environmental restructuring; modelling; and enablement (16), which were linked to 15 appropriate BCTs (14) (Table 2).

**Table 2.**
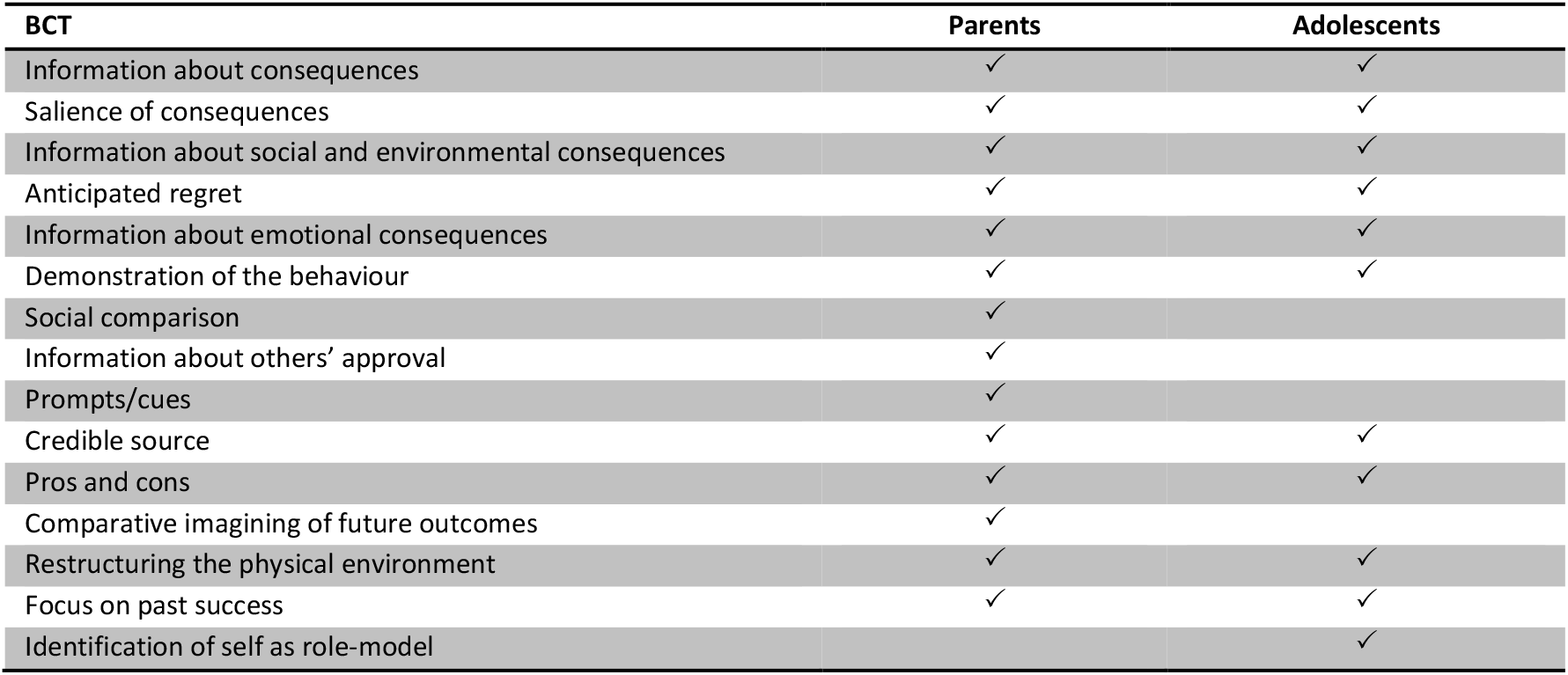
Behaviour Change Techniques (BCT) identified through qualitative research (11)

| BCT | Parents | Adolescents |
| --- | --- | --- |
| Information about consequences | ✓ | ✓ |
| Salience of consequences | ✓ | ✓ |
| Information about social and environmental consequences | ✓ | ✓ |
| Anticipated regret | ✓ | ✓ |
| Information about emotional consequences | ✓ | ✓ |
| Demonstration of the behaviour | ✓ | ✓ |
| Social comparison | ✓ |  |
| Information about others' approval | ✓ |  |
| Prompts/cues | ✓ |  |
| Credible source | ✓ | ✓ |
| Pros and cons | ✓ | ✓ |
| Comparative imagining of future outcomes | ✓ |  |
| Restructuring the physical environment | ✓ | ✓ |
| Focus on past success | ✓ | ✓ |
| Identification of self as role-model |  | ✓ |

Therefore, the aim of this study was to design, develop and evaluate a theory and evidence-based intervention to improve HPV and HPV vaccine knowledge, and intention to vaccinate, among parent-daughter dyads.

## Methods

Ethical approval was obtained from the Social Research Ethics Committee, University College Cork (Log 2019-26). An online behavioural intervention entitled *“Is the HPV vaccine for me?”* was developed. The six minute video was created using VideoScribe 3.3.1-1 software by Sparkol^®^, in consultation with a Technology-Enabled Learning Co-ordinator. A narrative approach was applied, mapping the adolescent HPV vaccine decision journey (BCT: demonstration of the behaviour; social comparison). It was narrated by the primary researcher and definitions and numerical information were complemented by graphical illustration. The information provided is evidence-based and theoretically-informed, bridging the knowledge gaps identified through previous research (11). It addresses the objectives outlined in Table 3, according to the identified BCTs. The video finishes with a reminder that the majority of girls in Ireland accept the HPV vaccine (BCT: information about social and environmental consequences; social comparison; information about others’ approval).

**Table 3.**
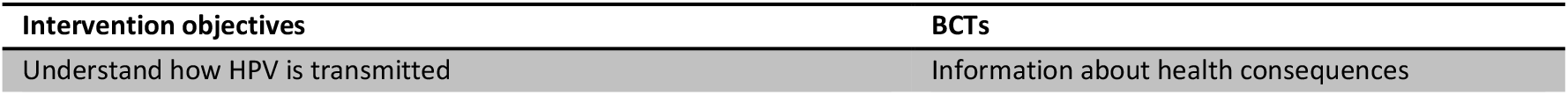
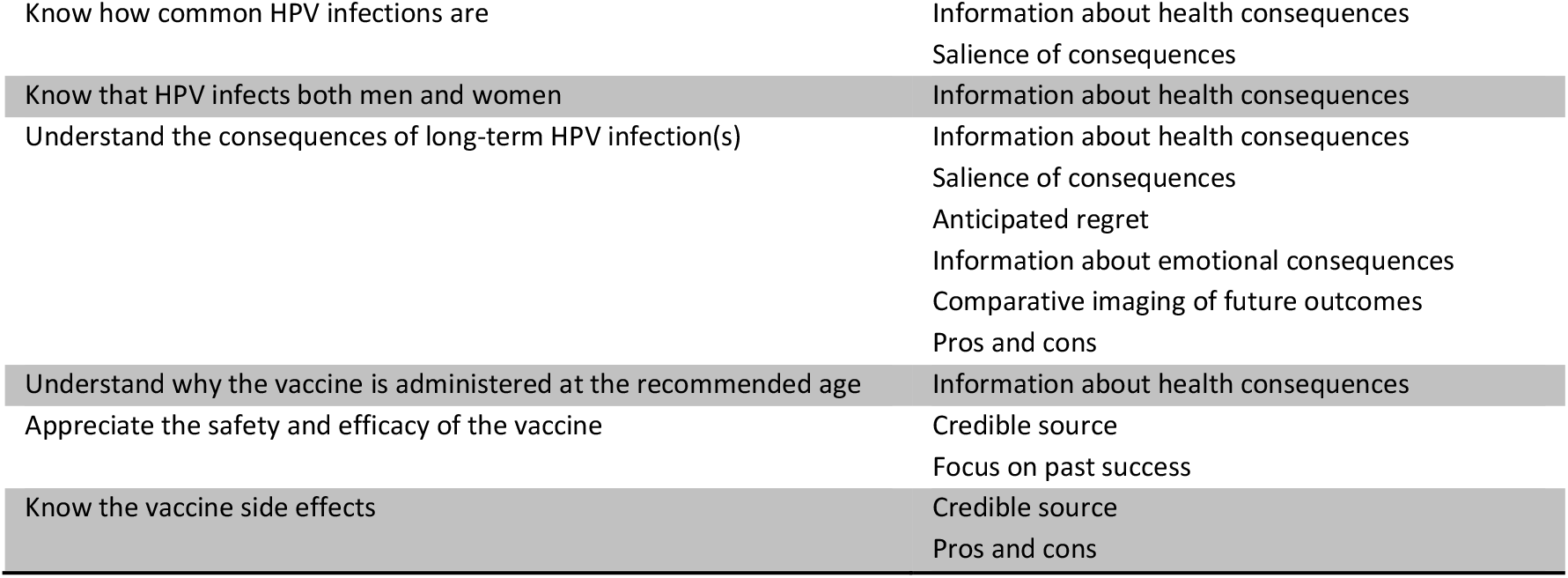
Intervention objectives and associated Behaviour Change Techniques (BCTs) (14)

A prospective randomised controlled feasibility trial (RCT), containing an intervention group who had access to the video and a control group who did not have access to the video, was conducted to evaluate the intervention in Cork, Ireland. Eligible participants were parent-adolescent dyads including a female adolescent, pre-HPV vaccination, in her final year of primary school (ISCED level 1) (17), typically aged 11-12 years. Recruitment took place over a six-week enrolment period from April to May 2019. A list of primary schools was compiled and stratified according to DEIS (Delivering Equality of Opportunity in Schools) status (18). The DEIS programme supports children who are at greatest risk of educational disadvantage (18). Using a purposive sampling strategy, school principals were contacted via email and/or telephone and provided with details of the trial. Schools interested in participating were then randomised by simple randomisation. The principals were asked to share study information, via email, with eligible participants. This email detailed trial information, expectations of participation, and instructions on accessing the trial material. Google Forms was used as the data collection platform, recording consent to participate and baseline characteristics of the parent: gender; age range; highest education level achieved; number of children (under 18 years); and vaccination status of children. A 10-item questionnaire was developed to evaluate participants’ baseline HPV and HPV vaccine knowledge and intention to vaccinate (Figure 1). Items assessed knowledge using a “True, False, Don’t know” format. These were identified as knowledge gaps during previous literature review and qualitative research (10, 11). The questionnaire was developed, edited and assessed for face and content validity but did not undergo external validation. A knowledge score was based upon correct responses to items, with one point being rewarded for each correct response obtained (range 0-10). No points were rewarded for “Don’t know” responses. Participants were also asked about their intention to accept the HPV vaccine (“Yes, No, Don’t know”). This question was not scored. At the outset of the study, (W0), all participants undertook baseline knowledge assessment. Those in the intervention group were immediately invited to view the video. Two weeks later (W2), all participants repeated the knowledge assessment. Those in the intervention group were asked whether the video had increased the likelihood of accepting the HPV vaccine (“Yes, No, Don’t know”). After completing knowledge assessment at W2, those in the control group were provided the opportunity to view the video. At W2, participants in the intervention group were asked the following questions: “Did you find this video interesting?” and “Did you find this video useful?” (“Yes, No, Don’t know”). In addition, feedback from participants was obtained through the provision of a free-text box. Data were analysed using IBM’s SPSS. Continuous variables were described by medians and IQRs (non-parametric data). Categorical variables were described by counts and percentages. Associations between categorical variables were investigated using Yates’ Continuity Chi-Square tests. Mann-Whitney U tests were used to investigate differences between groups for continuous variables. P values of <0.05 were considered statistically significant.

**Figure 1.**
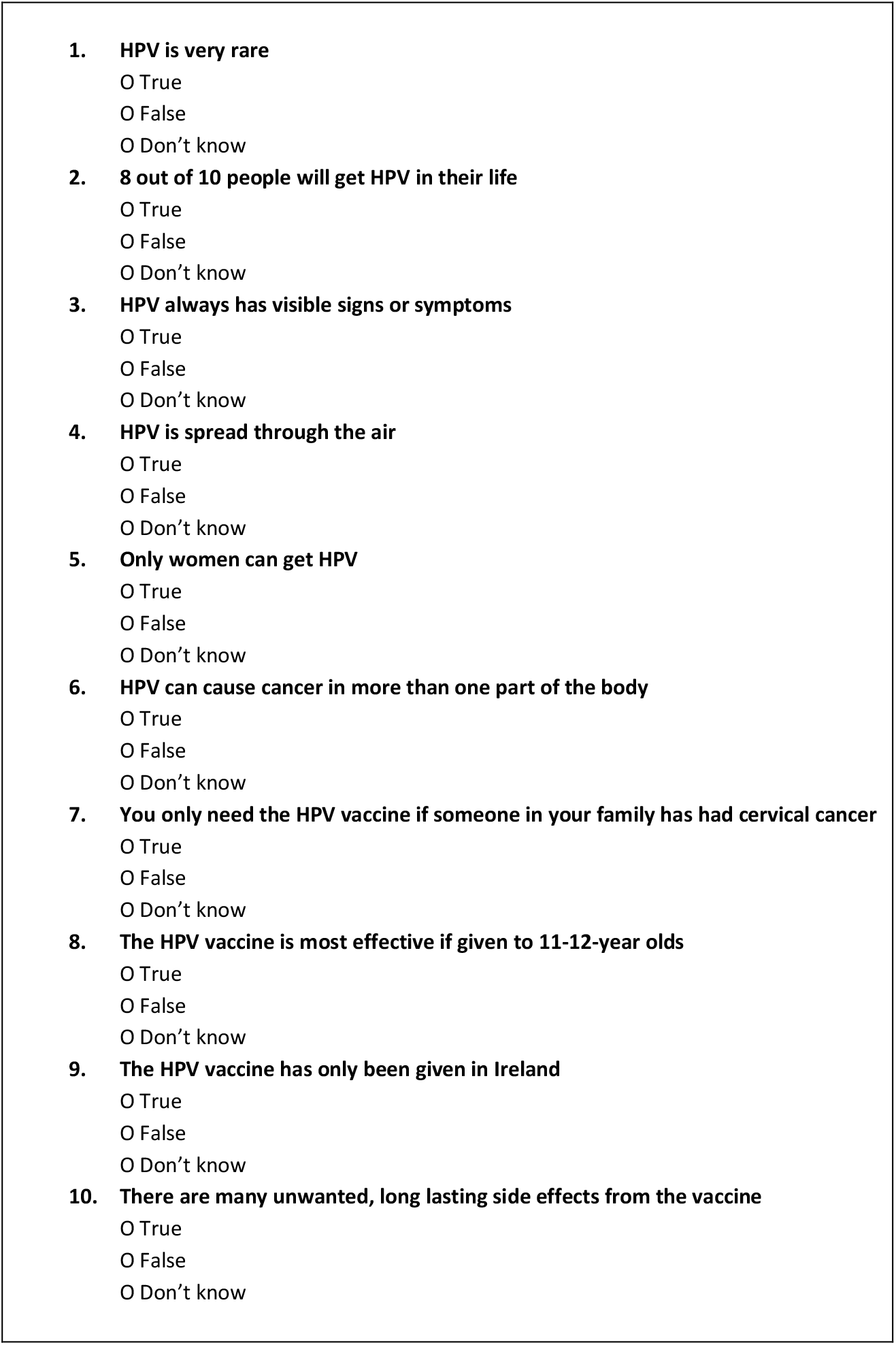
HPV and HPV vaccine knowledge questionnaire.

## Results

A total of 313 schools were invited to participate (n=37 DEIS, n=276 non-DEIS). Eleven schools agreed to participate (n=4 DEIS, n=7 non-DEIS) and were randomised (n=5 control, n=6 intervention). According to information provided by schools, 326 parent-daughter dyads were eligible to participate. A total of 49 dyads completed baseline knowledge assessment at W0 (n=24 control, n=25 intervention), and 35 dyads completed knowledge assessment at W2 (n=17 control, n=18 intervention). All participants were female and declared that, to the best of their knowledge, all children in their care were fully vaccinated. There were no statistically significant differences between the control and intervention groups based on DEIS status (χ^2^_yates_ (1)=0.013, p=0.909), age (χ^2^_yates_ (1)=3.463, p= 0.063), education (χ^2^_yates_ (1)=0.000, p=0.984), number of children (χ^2^_yates_ (1)=0.580, p=0.456) or intention to accept the HPV vaccine (χ^2^_yates_ (1)=0.021, p=0.884).

At W0, the overall median (IQR) baseline knowledge score was 5 (4, 6). There was no statistically significant difference in baseline knowledge assessment scores between control and intervention groups, (U=292, Z=−0.163, p=0.870) (Figure 2). Just over half (51%) of the participants indicated that they intended to accept the HPV vaccine, while the remaining 49% remained undecided.

**Figure 2.**
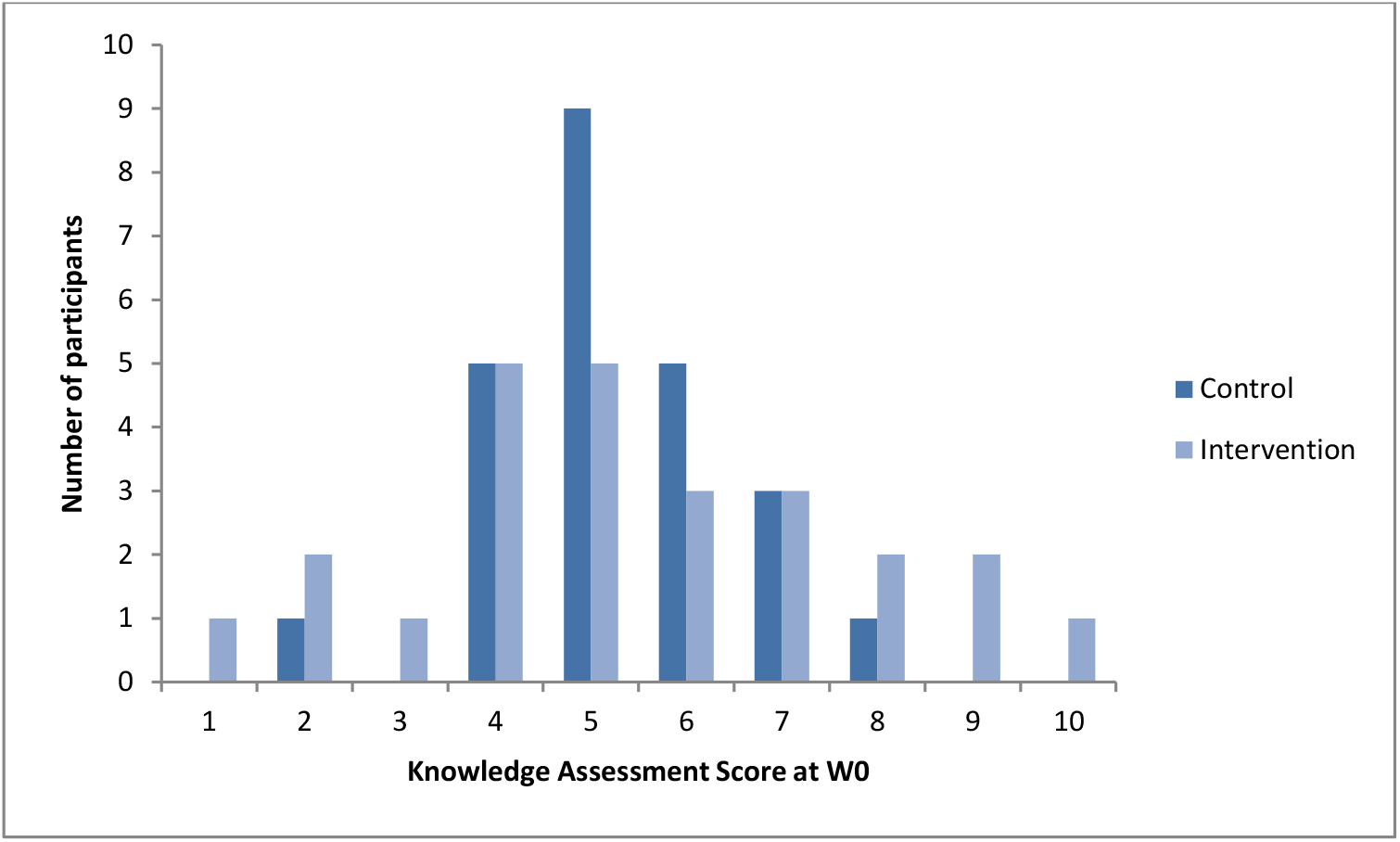
Knowledge Assessment Scores at W0, comparing Control and Intervention groups.

At W2, there was a statistically significant difference in knowledge assessment scores between control and intervention groups (U=1.5, Z=−5.065, p<0.01) (Figure 3). When asked whether this video had increased the likelihood of accepting the HPV vaccine, 88% indicated that it had, 4% indicated that it had not and 8% were unsure. All intervention participants found the video interesting, while 96% found it useful.

**Figure 3.**
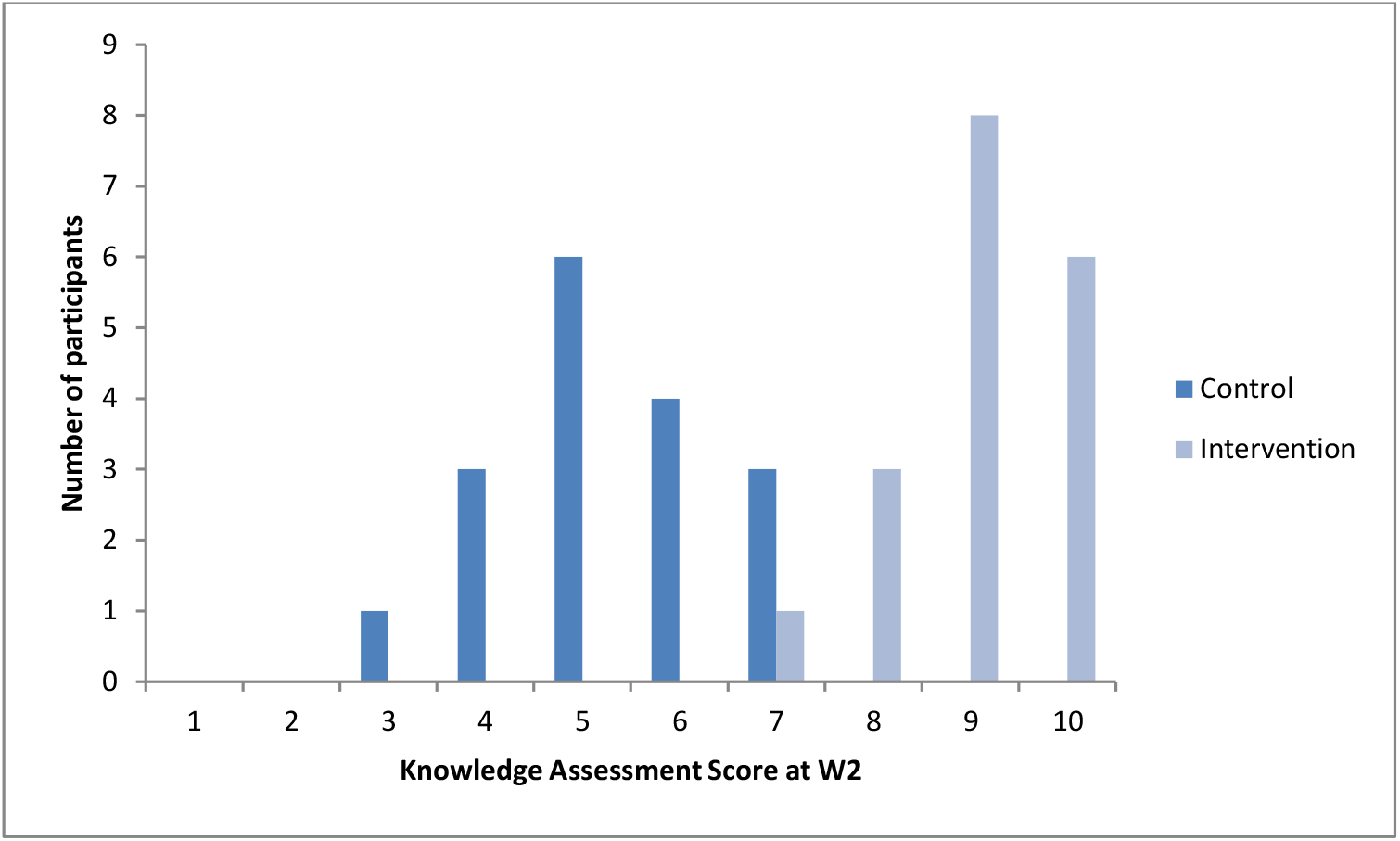
Knowledge Assessment Scores at W2, comparing Control and Intervention groups.

## Discussion

The purpose of this study was to design, develop and evaluate the feasibility of a theory, and evidence-based intervention to improve knowledge about HPV and HPV vaccines and intention to vaccinate among parent-daughter dyads. It was intended that targeting these dyads would promote open dialogue between parent-daughter pairs, leading to a scenario where the adolescent was involved and participated in the vaccine decision. The chosen mode of delivery was an online video. Digital media has several advantages: videos can be entertaining; their medium is familiar; and they may be designed as a “takeaway tool” that permits more independent application, at the viewer’s own pace (19). We found that this educational intervention significantly increased knowledge for participants who viewed the video. Secondly we found that this video increased the likelihood of accepting the HPV vaccine for the majority of participants. Therefore this study provides initial “proof of concept” that an educational intervention designed from a solid foundation of behaviour change theory can positively influence HPV vaccine decision-making.

While several interventions have been designed to address HPV vaccine hesitancy (6, 7, 20, 21), there are no published examples, using the BCW to develop a *de novo* online intervention, targeted at parent-daughter dyads. The intervention was evaluated using the APEASE criteria: a set of criteria used to make context-based decisions on intervention content and delivery consisting of affordability, practicability, effectiveness and cost-effectiveness, acceptability, side-effects/safety and equity considerations (13). While affordability is difficult to quantify in this case, this video-based intervention was created in consultation with a Technology-Enabled Learning Co-ordinator, with minimal financial input. This video was hosted free of charge on a YouTube^®^ platform (unlisted). This intervention could be adopted in its current form with no further financial investment. It is practicable in its mode of delivery: the video is hosted online and the internet is used as a platform to disseminate the content to the target population. According to data collected in 2019 by Ireland’s Central Statistics Office (CSO), an estimated 91% of Irish households had access to the internet at home (22), with 57% of individuals seeking health-related information online (23). More people are accessing internet-based content by following links on social media than through direct searches (24). Social media statistics from June 2018 indicate that up to 66% of Irish individuals (over 15 years) are using social networking sites (e.g. Facebook^®^, Instagram^®^, LinkedIn^®^, Twitter^®^) (25). It has been demonstrated that information shared via social media results in greater knowledge transfer than when shared via pamphlets (26). In addition, it has been postulated that social media has direct public health relevance because social networks could have an important positive influence on health behaviours and outcomes (27, 28). Therefore, social media platforms have the potential to effectively increase knowledge and facilitate behaviour change. The intervention described in this study is readily amenable to dissemination on social media platforms.

In this small feasibility study, the intervention was shown to be effective, with a statistically significant increase in knowledge assessment scores. At the outset, 49% of participants were undecided about their vaccine decision. On completion of the study, 88% of participants indicated that the video had increased the likelihood of accepting the vaccine. While this was a positive finding, it must be acknowledged that intention alone does not necessarily predict future vaccine uptake (29), a disparity known as the intention-behaviour gap (30). A variety of strategies have been suggested to bridge this gap: keeping these favourable immunisation intentions in mind through reminders, prompts and cues; and reducing barriers through logistics and heuristics (31).

There were no statistically significant differences between groups according to DEIS status, age range, education, number of children, child vaccination status, and intention to accept the HPV vaccine. All parent participants were female. However, such a gender imbalance is not unusual. Research has demonstrated that the female care-giver is more likely to participate in clinical research (32), and is often the primary healthcare decision-maker for the family (33).

While it would have been desirable to evaluate the impact of this intervention on actual vaccine uptake, this was not a primary objective of the study. Instead, the favourable immunisation intentions generated by the intervention could be kept in mind through repeated dissemination of the video (e.g. online via social media or on television screens located at healthcare facilities such as GP surgeries and pharmacies). Due to the affordability, practicability and effectiveness of the intervention, it was determined to be cost-effective.

Acceptability has become a key consideration in the design, evaluation and implementation of healthcare interventions and is a necessary condition for effectiveness (34). The acceptability of this intervention was evaluated: all of those asked found the video interesting, while 96% found it useful. However, only 3.5% of invited schools consented to participate. While an effort was made to understand the reasons underpinning their lack of participation, the majority of contacted schools were non-responsive. Of those eligible to complete the knowledge assessment at W0, only 15% did so and this was further reduced to 10.7% at W2. This decline in response rate between phases is frequently observed (35). However, a higher response rate was expected due to the personal relevance of the research topic (36). This intervention was delivered in May i.e. four months before the vaccine would be offered to the participants, to permit timely provision of vaccine information (37). It is possible that the time-lag between information delivery and actual vaccination date was too long, and participants were not yet prepared to consider their vaccine decision and thus were not personally invested in intervention content.

An intervention may be effective and practicable, but have unwanted side-effects or unintended consequences (13). Research has shown that information provision regarding vaccine safety and efficacy can cause unpredictable effects on vaccination uptake and may even increase such concerns (38, 39). A free-text box was provided in this study and no participants reported any such concerns. However, the potential for such an occurrence if the intervention were to be scaled up should be considered.

The use of video as a mode of delivery facilitates equity, as it provides standardised content across learners and has been shown to be effective among viewers of lower literacy levels (40). However, the impact of the ‘digital divide’ on the implementation of an online intervention must be considered. A proportion of the Irish population (11%) do not have internet access at home with 30% of these reporting that lack of skills hampered their internet access (23). Parents (and adolescents) currently receive vaccine information in written format, (i.e. pamphlets and patient information leaflets (PIL)) and additionally, the website of the National Immunisation Office (NIO) is signposted, providing further information in a variety of formats. The intervention described here is not intended to replace such material, but rather to support and complement it, providing information and promoting behaviour change in a manner that is independent of the Government.

## Conclusions

A video-based online behavioural intervention was associated with improved HPV (and HPV vaccine) knowledge, and intention to vaccinate, among parent-daughter dyads. The intervention was found to be affordable, practicable, effective (cost-effective), acceptable, safe, and equitable, in this feasibility study. However, it is important to acknowledge that vaccination is highly context specific (41). Therefore the impact of this intervention will need to be evaluated in alternative contexts. In addition, from September 2019, male adolescents are included in the HPV vaccination programme (1). An assessment of the impact of this intervention in parent-son dyads is required, making alterations as required, and supplementing with further qualitative research, if indicated. Should this intervention demonstrate efficacy across multiple contexts, a national dissemination of *“Is the HPV vaccine for me?”* should be launched.

## Acknowledgements

The authors wish to acknowledge the valuable contribution of Mr Patrick Kiely, Technology-Enabled-Learning Co-ordinator in the Office of the Vice-President for Teaching and Learning, University College Cork, the principals of the participating schools, and the study participants, to this research.

